# Learning and recognition of tactile temporal sequences by mice and humans

**DOI:** 10.1101/122887

**Authors:** Michael Bale, Malamati Bitzidou, Anna Pitas, Leonie Brebner, Lina Khazim, Stavros Anagnou, Caitlin Stevenson, Miguel Maravall

## Abstract

The world around us is replete with stimuli that unfold over time. When we hear an auditory stream like music or speech or scan a texture with our fingertip, physical features in the stimulus are concatenated in a particular order, and this temporal patterning is critical to interpreting the stimulus. To explore the capacity of mice and humans to learn tactile sequences, we developed a task in which subjects had to recognise a continuous modulated noise sequence delivered to whiskers or fingertips, defined by its temporal patterning over hundreds of milliseconds. GO and NO-GO sequences differed only in that the order of their constituent noise modulation segments was temporally scrambled. Both mice and humans efficiently performed tactile sequence learning. Mouse performance relied mainly on detecting relative changes in noise amplitude over time, whereas humans appeared to have access to more cues, including the duration of noise modulation segments.

## Introduction

To make sense of the world around us, the brain must integrate sensory patterns and sequences over time and assign them meaning. Signals in our environment unfold over time and can only be interpreted by decoding their temporal patterning. The ability to do so underpins much of our sensory experience – for example, it is central to recognising a favourite melody or a passage of speech [1]. As first proposed over 60 years ago [2], sequence processing provides a model for investigating how neuronal circuits give rise to object perception and recognition, a central goal of neuroscience [3, 4].

In tactile sensation, fast sensory events, such as fluctuations in the forces acting on a whisker follicle, are encoded faithfully and with high temporal precision [5-11]. Exploring an object by scanning with fingertips or whiskers generates a series of tactile events concatenated over time [12-17]. Recognising the object as a whole – its texture, shape or size – requires integrating over these events, with the relevant timescales varying from tens of milliseconds (ms) to seconds.

Introspection suggests that meaningful auditory sequences, such as those in speech or music, can be learned quickly and robustly. We wondered whether similarly effective sequence learning and recognition occurs in tactile sensory systems. We further wished to explore the cues that could underlie capacities for tactile sequence recognition.

To address these issues, we developed a new experimental design for testing sequence discrimination in mice and humans. Participants learn to distinguish a target stimulus sequence, constructed from an underlying noise waveform, from other stimuli that differ only in their temporal patterning over hundreds of milliseconds. Our results demonstrate efficient learning of tactile sequences both in mice and in humans. This behaviour provides an assay for exploring the neuronal circuit mechanisms that underpin recognition of temporally patterned stimuli.

## Materials and Methods

### Surgical procedures

All procedures were carried out in accordance with institutional, national (Spain and United Kingdom) and international (European Union directive 2010/63/EU) regulations for the care and use of animals in research. Details of head bar implantation surgery have been described elsewhere [7, 18]. Briefly, under aseptic conditions, mice (male, total n = 32, 6-9 week old) were anaesthetised using 1.5-2.5% isoflurane in O**2** and placed into a stereotaxic apparatus (Narishige, Japan) with ear bars previously coated with EMLA cream. We monitored anaesthetic depth by checking spinal reflexes and breathing rates. Body temperature was maintained at 37°C using a homeothermic heating pad. Eyes were treated with ophthalmic gel (Viscotears Liquid Gel, Novartis, Switzerland) and the entire scalp was washed with povidone-iodine solution. An area of skin was removed (an oval of 15 mm × 10 mm in the sagittal plane) such that all skull landmarks were visible and sufficient skull was accessible to securely fix a titanium or stainless steel head bar. The exposed periosteum was removed and the bone was washed using saline solution. The bone was dried and then scraped using a scalpel blade to aid bonding of glue. Cyanoacrylic glue (Vetbond, 3M, USA) was applied to bind skin edges to the skull and as a thin layer across the exposed skull to aid bonding to the dental acrylic. A custom titanium or stainless steel head bar (dimensions 22.3 × 3.2 × 1.3 mm; design by Karel Svoboda, Janelia Farm Research Campus, Howard Hughes Medical Institute) [18] was placed directly onto the wet glue centred just posterior to lambda. Once dry, we fixed the head bar firmly in place by applying dental acrylic (Lang Dental, USA) to the head bar (on top and behind) and the skull (anterior). Mice were given buprenorphine (0.5 mg/kg, I.P.) and further EMLA cream to the paws and ears. Once the acrylic was set, anaesthesia was turned off. Animals were housed individually on a reverse 50:50 light-dark (LD) cycle and allowed to recover for one week post-surgery.

### Head fixation and water delivery

Mice were trained using a shaping procedure to freely enter a head fixation device (Figure 1A). We used two device designs. One design consisted of an acrylic tube (32 mm internal diameter) with its head end cut to enable access to implanted head bars. The tube was placed on Parafilm or a rubber glove and clamped into a v-shape groove. This support acted to stabilise the tube, collect faeces and prevent mice from grasping stimulus apparatus and the lickport. The second design consisted of a platform with a custom-made treadmill on which mice could locomote freely (design by Leopoldo Petreanu, Champalimaud Centre for the Unknown). A mesh was fixed over the treadmill to surround the mouse’s body, allowing the animal to feel comfortably enclosed rather than exposed. The ends of the head bars were inserted into grooves on two head fixation clamps and tightened using thumbscrews. The head fixation set-up was adapted from [18, 19].

**Figure 1.**
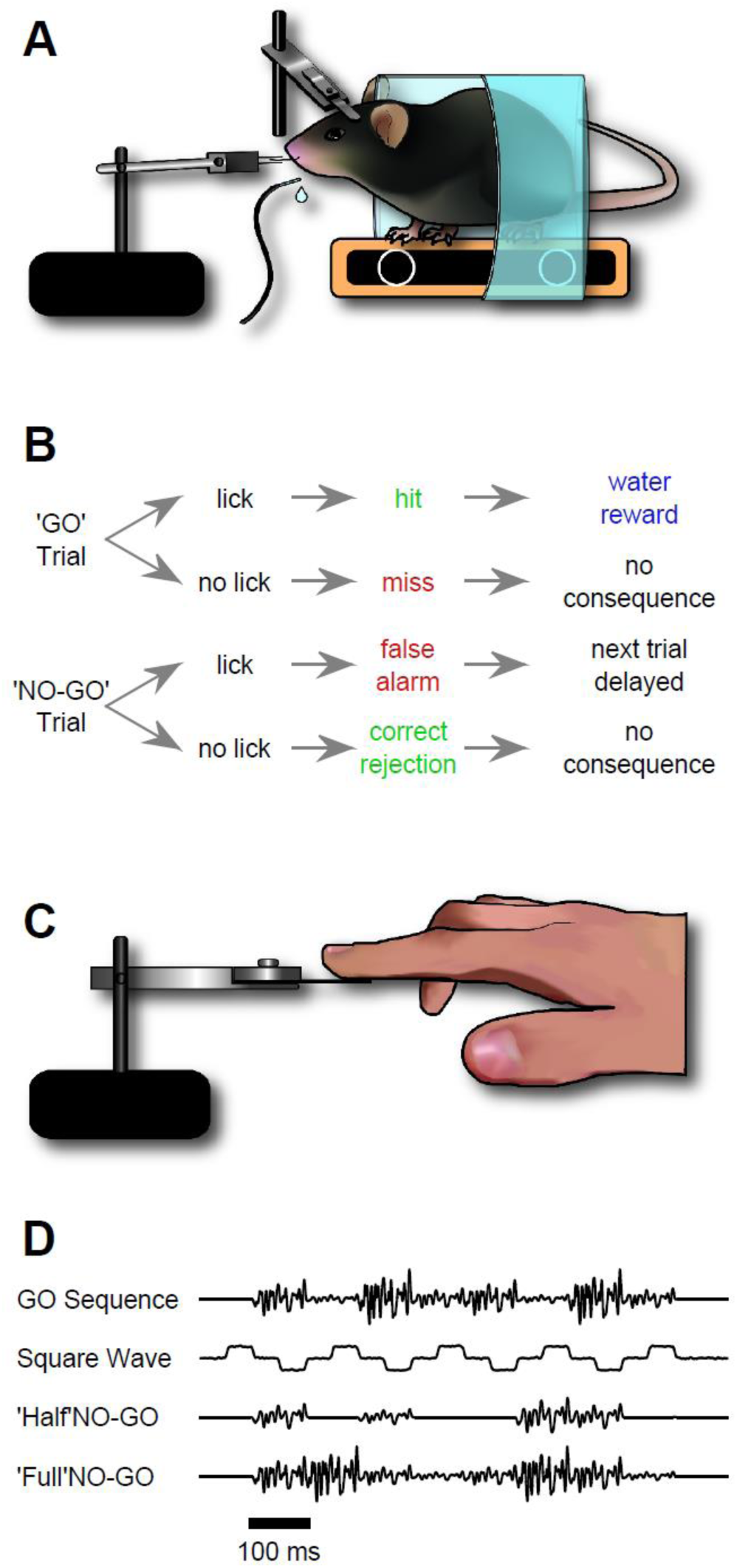
Design of sequence recognition task for mice and humans. A. Illustration of the treadmill-based behavioural and stimulus delivery setup for head-fixed mice. B. Block diagram representing the structure of the GO/NO-GO paradigm. On GO trials, a mouse licking the water spout within the response period (hit) was rewarded with a water droplet. If the mouse licked on a NO-GO trial (false alarm), the next trial was delayed by 2-5 s. C. Illustration of the stimulus delivery setup for human experiments. D. Stimulus sequences for discrimination and intermediate ‘shaping’ sequences.

Water was available to mice via a spout made from a blunted gauge 13 syringe needle. Water delivery was controlled via a solenoid valve (LDHA1233215H, The Lee Company, France). The acrylic tube or head bar holder was lined with aluminium foil. Terminals from an A/D input of a signal processor (RP2.1, TDT, USA) were then connected to the water spout and the foil. Tongue contacts with the lick port created brief elevations in voltage consistent with lick durations [20].

### Water restriction

To motivate mice to learn and perform the task we employed a water restriction protocol [18] and made water available as a reward during the task. Mice cope better with water control than food control [21]. Unless rodents are motivated by fluid or food control, they can fail to learn even simple sensory tasks [22] and perform too few daily trials for data collection to be satisfactory. We verified that mice were not motivated by sugary treats alone (Lucozade and chocolate milk). We observed a mild increase in motivation when mice were given sunflower seeds before tasks.

Mouse water intake was regulated so that animals were motivated to perform at around 75% success rate for 200 or more trials per session under our conditions (45-55% humidity, 23°C and atmospheric pressure; reverse 50:50 LD cycle), while remaining active and healthy. This was achieved with two different schedules, depending on the institution where the experiment took place. In one schedule (Instituto de Neurociencias), we titrated down water availability to the amount required for mice to maintain >75% of initial body mass in the short term and gradually increase body mass in the long term (0.5 ml daily including experimental water rewards collected during the session, 7 days a week). In the other schedule (University of Sussex), mice were restricted to 50% of their average free water intake but given free access to water for a finite period during the dark phase of their LD cycle. Body weight (mass) was monitored throughout the study, and we measured experimental reward water intake by weighing mice before and after the daily behaviour session together with collected faeces. For both schedules, mice initially lost weight but then gradually increased body mass over the course of the experiment. Sensory discrimination training began after 9 days on water control.

### Animal handling and training

We initiated water control one week after head bar implantation, and began to handle animals daily. On days 1 and 2 animals were introduced to the experimenter. On days 3 and 4 animals were introduced to the head fixation device. On days 5 and 6 mice received water via a syringe only when inside the device (but not head-fixed). On days 7 and 8 animals were given a sunflower seed and after ingestion were head-fixed and given water via a syringe. Animals became accustomed to head fixation and expected to receive water from the spout situated in front of their head. On day 9, under light isoflurane anaesthesia (1-2%) all whiskers apart from C2 were trimmed bilaterally. At least 30 minutes later mice began the task. Mice performed a single daily training session. Animals were trained in the dark; illumination, if necessary, was provided by a red lamp.

### Stimulus delivery and design

Our aim was to develop a task whereby tactile sequences delivered to the animal could only be distinguished by discriminating their temporal patterning. Careful control of stimulation patterns was therefore required. To achieve this we delivered controlled stimuli, which animals needed to sense by operating in a “receptive” mode rather than by active whisking [23]. In this design, whiskers were inserted into a small tube. Stimulus sequences were generated as filtered noise vibrations, such that whisker stimulation was continuous during a trial (Figure 1D). We thus avoided temporally isolated discrete movements that could have initiated whisking or confused the animal as to the start, content and ending of the temporal pattern. Upon head fixation at the start of a session, the left C2 whisker was inserted into a snugly fitting tube (pulled 1 ml plastic syringe) glued to a piezoelectric actuator wafer (PL127.11, Physik Instrumente, Germany). The wafer was mounted vertically and motion was rostrocaudal. In some experiments, a different method to deliver stimuli was required: a metallic 10 mm^2^ mesh grid was glued to the end of an actuator to enable multiple whisker stimulation and allow quick transition between experiments, as in Figure 2B,C.

**Figure 2.**
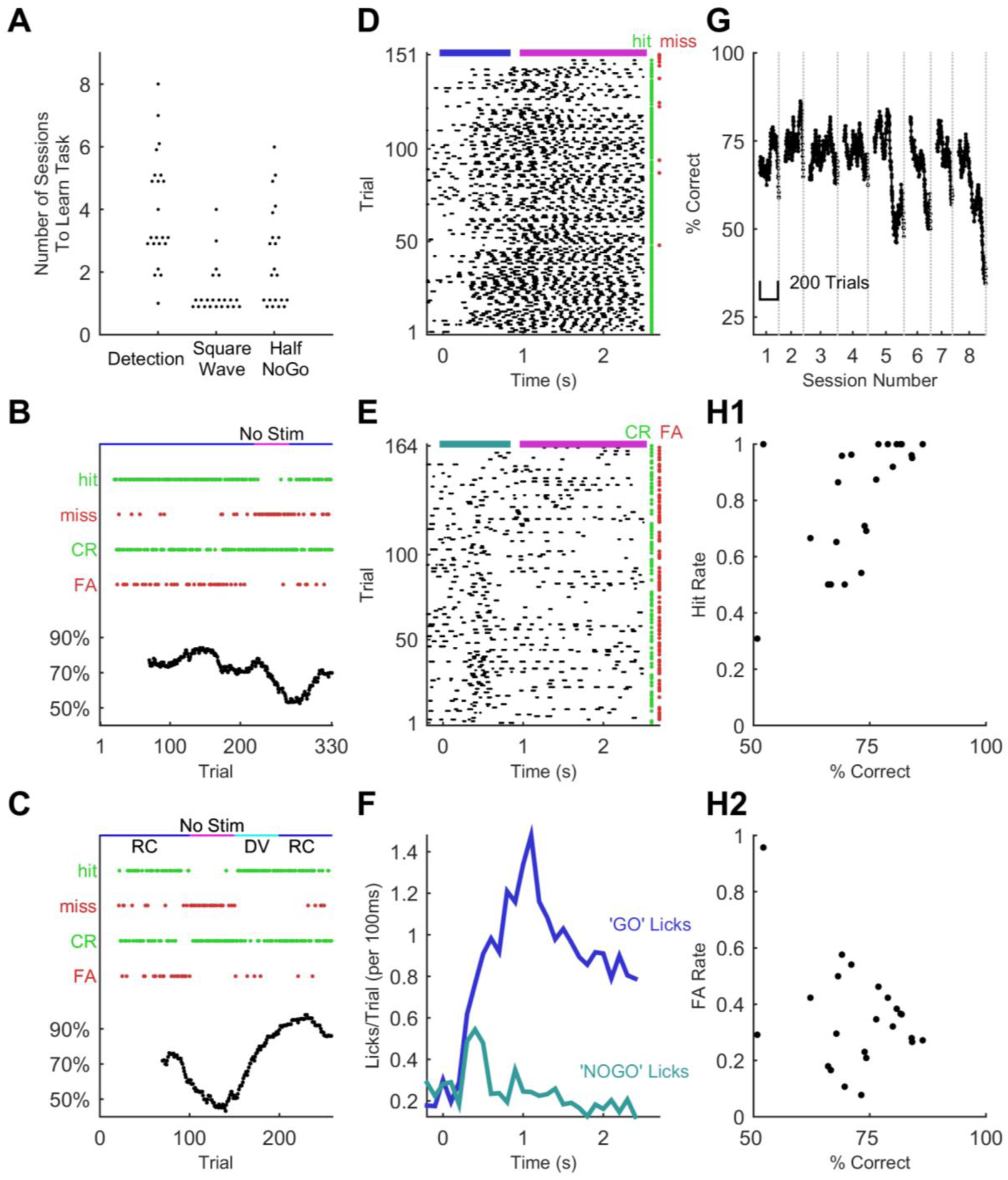
Sequence learning performance in mice. A. Number of training sessions needed to learn different training stages (75% performance criterion). Each dot, one mouse. Dots are jittered for visualisation. B. Performance metrics for an example training session (discrimination of the GO sequence from square wave). Correct trials are green dots, incorrect are red dots. Performance quantified as % correct over a 50-trial moving window (bottom). Whisker dependency on the task was verified by removing the actuator at trial 220 (fuchsia bar) and reinserting (blue bar) at trial 270. Apparent delay in performance is caused by 50-trial averaging. C. Performance for an example session with stimulus rotation (GO sequence versus square wave discrimination). Main symbols as for B. Stimuli were delivered, as normal, first in the rostro-caudal axis (RC; blue bar) but following a brief period of stimulator removal (fuchsia bar), in the dorso-ventral (DV; cyan) axis for 50 trials. Stimulation then returned to RC for the remainder of the session. D. Lick time raster plot for GO trials in an example session on the final stage of training (GO vs ‘full’ NO-GO). Licks are referenced to the start of the sequence, which lasts 800 ms (blue bar). The response period opens at 1 s (purple bar). Water was only available in the response period. Hit and miss trials are green and red dots, respectively. E. Lick time raster plot for NO-GO trials of same session as D. Grey-blue bar, sequence. Green dots denote correct rejection trials, red dots false alarm trials. F. Smoothed average of lick signals per trial from D and E. Blue, GO trials; grey-blue, NO-GO. G. Performance across 8 successive behavioural sessions for one mouse. Performance averaged over a 50-trial moving window. H. Hit and false alarm rates as a function of % correct. Each dot, one session on GO vs full NO-GO. The same sessions are depicted in H1 and H2.

Stimulus sequences were constructed in Matlab (Mathworks, USA) and played via a signal processor (RP2.1, TDT, USA) controlled with code custom-written in ActiveX software (TDT). The GO sequence lasted 800 ms and consisted of 8 consecutive “syllables”, where each syllable was a 100 ms segment constructed from white noise with one of 4 amplitude levels (Figure 1D). We constructed the sequence as follows: (1) we created a 100 ms white noise snippet generated at a sampling rate of 12207 Hz (in Matlab), (2) stitched 8 snippets together, (3) multiplied the resulting chain of repeated white noise snippets by an amplitude modulation envelope, (4) convolved this sequence with a Gaussian waveform (SD 1.64 ms) to implement frequency filtering, and (5) normalised the sequence to match the dynamic range of the piezoelectric actuator. In the resulting GO sequence, constituent syllables differed in amplitude: the pattern of noise amplitude modulation was [3 1 4 2 3 1 4 2], with 1 being the lowest amplitude level and 4 the highest. The NO-GO sequence in the full version of the task contained the exact same syllables but in a scrambled order (Figure 1D), specifically [3 4 2 1 2 4 3 1]. The target and non-target sequences were therefore identical for the initial 100 ms. Further sequences were created to aid learning and to explore the nature of recognition, as detailed in Results.

### Task control and analysis

We trained mice to respond to the GO sequence by licking a spout to receive a water reward (1-2 μl). On presentation of the NO-GO sequence mice were trained not to lick (Figure 1B). The trial began with the ‘stimulation period’ (0.8 s) where the sequence was delivered to the whisker. At the end of the stimulation period followed a ‘response period’ (1.5 s) where mice must lick or refrain depending on the stimulus sequence. Following the GO sequence, if mice licked during the response period (a hit trial) they received a water reward; if they failed to lick (a miss trial) the next trial began as normal. Following a NO-GO sequence, if mice correctly withheld licking during the response period (a correct rejection trial) the next trial began as normal; if they licked (a false alarm trial) the next trial was delayed by 2-5 s. Trial parameters were defined in Matlab using a custom made GUI and then loaded to the RP2.1 signal processor. Trial outcomes were recorded in Matlab using custom-written code.

Several related measures can be used to quantify performance, including overall percentage of correct trials, hit rate and false positive rate, and d’ [22, 24]. Here we present results mostly as percentage of correct trials measured over a 50-trial sliding window during the course of a session. To calibrate this performance measure in terms of statistical significance level, we shuffled stimulus identity and behavioural response (lick/no lick) on a trial by trial basis for each individual session in a test data set of 104 sessions (n = 7 animals; shuffling repeated 10000 times per session). Performing shuffling separately for each session allowed us to control for variations in overall lick rate from animal to animal and during the course of training. For all sessions in the test data set, the probability of achieving 75% correct performance given a random relationship between stimulus and responses was lower than p = 0.001; the probability of achieving 70% correct given such a relationship was under p = 0.015.

During training, we routinely varied the proportion of GO and NO-GO trials during a session in order to aid learning and keep animals motivated (e.g. the fraction of GO trials could temporarily increase). This could lead to a misleading value of the performance measure. For example, consider a randomly performing mouse that licked on 90% of trials. In a hypothetical 50 trial period with 40 GO and 10 NO-GO trials, it would reach a 90% hit rate on GO trials and a 90% false alarm rate on NO-GO trials. Overall performance would then be 74% correct (= 0.8 × 90% + 0.2 × 10%), despite the mouse performing at chance with no differentiation between GO and NO-GO stimuli. To correct for this, we rebalanced the percentage correct measure so that GO and NO-GO trials are set to have equal weight. This rebalanced measure reports the above hypothetical example as 50% correct (= 0.5 × 90% + 0.5 × 10%).

### Human experiments

Human experiments were conducted and underwent ethical review at the University of Sussex. In total, 59 participants were recruited and gave informed consent. In the human counterpart of the experimental design, the basic GO and NO-GO stimulus waveforms described above (Figure 1D) were left unchanged. Further waveforms were added in order to aid and test learning as described in Results. Stimuli were loaded to the RP2.1 signal processor and delivered via a piezoelectric wafer identical to that used for whisker stimulation, but with a plastic plate glued on (polyethylene terephthalate; 20 × 10 × 1 mm). The wafer stimulator assembly was supported by a platform incorporating a cushioned armrest. Participants were asked to place one fingertip lightly on the plate’s surface (Figure 1C). The wafer was placed horizontally and vibrations were vertical. A small box containing a button was placed on the same table as the platform, in a position allowing participants to comfortably press the button with their free hand whenever a target stimulus was felt. GO and NO-GO stimulus trials were randomly interleaved. Experiments were conducted with no explicit instruction as to the identity of the target stimulus; instead, participants were asked to press the button whenever they identified a stimulus that felt familiar, more frequent or “special” than others. Participants had to decide by themselves which stimulus constituted the target. Feedback upon correct trials, provided in the form of a “Correct” sign appearing on a computer screen, was given to a subset of participants to more closely mirror the experimental design used with mice. We compared performance with and without feedback: performance was no higher for the participants trained with feedback, so results were pooled together (p = 0.99, Wilcoxon rank sum test, n = 15 participants without feedback and n = 44 with feedback).

## Results

### Achieving sequence recognition by mice

We sought to train mice to recognise a target stimulation sequence delivered to their whiskers. Our aim was for mice to distinguish the target sequence based on the order in which its elements appeared. To this end, mice were trained to distinguish between initially meaningless GO and NO-GO sequences built from series of identical “syllables”, with the sequences differing only in that syllables were scrambled in time over hundreds of milliseconds (each individual syllable lasting 100 ms, for a total of 8 syllables; Figure 1D). The initial syllable was identical across GO and NO-GO sequences in order to avoid providing a stimulus onset cue (Figure 1D).

Mice (n = 22) were trained to associate the GO stimulus with a water reward by making water available when the GO stimulus was delivered; on the first few days of training, no other whisker stimuli were given, so that mice effectively learned to detect whisker stimulation. As soon as animals demonstrated detection (75% correct detection trials), we introduced an initial NO-GO sequence. To make this stage easier, this initial NO-GO sequence consisted of a square wave riding upon low amplitude noise, distinctly different from the GO sequence (Figure 1D). Mice quickly learned to distinguish the square wave stimulus from the GO waveform (75% correct; within 4 sessions; Figure 2A). They were immediately moved to the next stage of training to avoid creating an artefactual generalized association between “noisy” stimuli (as opposed to square waves) and water availability. In the following —more demanding— stage, the NO-GO sequence consisted of a scrambled GO sequence with half (4 of 8) syllables knocked out (Figure 1D).

Mice accomplished each stage of training within a few days (Figure 2A), performing approximately 200-300 trials per daily session (mean 249 trials; SD 71 trials; total n = 456 sessions in 22 mice). During this process, recognition of the GO sequence was mediated by the animal’s whiskers: performance fell to chance level upon removing the whiskers from the moving stimulator (Figure 2B). Performance recognising the GO sequence was robust against variations in how the sequence was presented: daily changes in the tube’s positioning relative to the stimulated whisker did not noticeably affect performance. To test this invariance more specifically, in a subset of experiments mice were trained on a multi-whisker version of the task where whiskers (left untrimmed) were inserted into a wire mesh attached to the piezo actuator. Whiskers were first removed from the stimulator mesh; then, after a period of trials with stimulator movement but no whisker stimulation —during which performance dropped to chance level—, the actuator was rotated 90° and whiskers reinserted into the mesh. Reinsertion and stimulator rotation changed the identity and set-point of whiskers being stimulated as well as the direction of stimulation. Yet performance recovered to the level reached before whisker removal (Figure 2C; repeated for n = 4 mice; p = 0.44; Wilcoxon signed rank test). Sequence recognition therefore transferred across different stimulation configurations.

In the final stage of training, the NO-GO sequence comprised identical syllables to the full GO sequence but scrambled in time: that is, syllable ordering changed (Figure 1D). Mice that underwent training up to this final stage performed beyond 70% on at least one session (mean 72.8%, SD 9.4%; n = 5 out of 6 mice). Animals could maintain their performance over several days, albeit with fluctuations (Figure 2G), again despite day-to-day variability in how the whisker was attached to the stimulator. On every stage of training, improvements in performance occurred mainly through learning to withhold impulsive false alarm responses (licks) (Figure 2D-F,H), in common with other discrimination tasks in mice [18, 25]. Thus, high performance was associated with low false alarm rates (Figure 2H). These experiments show that mice learned to distinguish whisker-mediated stimuli that differed from others only in their temporal patterning over hundreds of milliseconds.

### Rapid learning of noisy sequences by humans

We tested whether humans could also learn to recognise the same temporally patterned tactile noise stimuli, delivering the patterns through an actuator applied to a fingertip. Participants were asked to indicate recognition of the target sequence by pressing a button. They first underwent a training session in which the GO target sequence was interleaved with a series of non-target stimuli. Early in the session, the non-target patterns were easily distinguishable from the GO pattern; later, these clearly distinct patterns were replaced by NO-GO sequences that differed from the GO target only in that their constituent syllables were scrambled, as in the final stage of mouse training. Human participants quickly improved their performance over the course of this session, typically converging to a steady level of performance despite the increase in difficulty during the session (Figure 3A). This indicated fast learning of the GO sequence. Performance as tested after the end of this learning phase was maintained in a later session separated by at least one week, suggesting remarkable robustness (Figure 3B; retest performance was actually higher, although the difference did not reach significance; p = 0.0547; n = 11 participants; Wilcoxon signed rank test).

**Figure 3.**
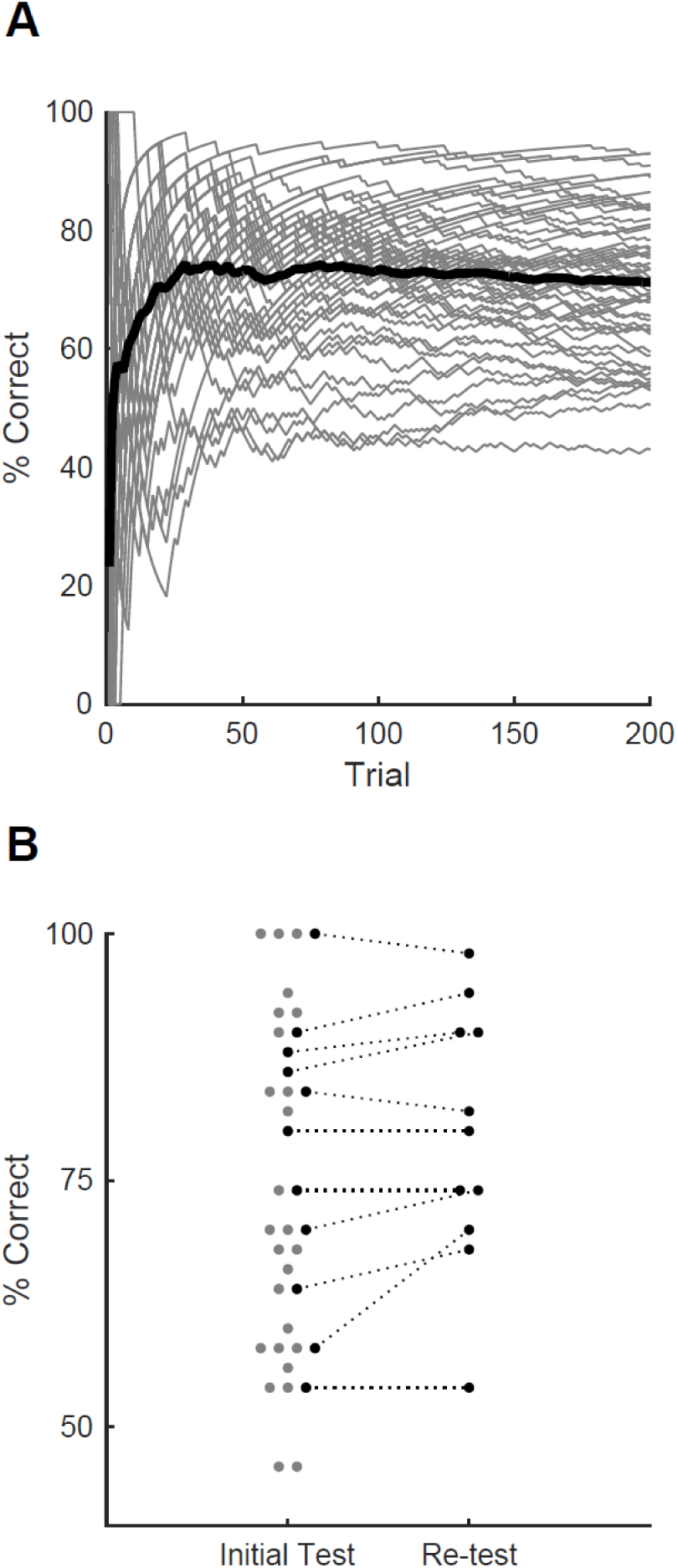
Sequence learning performance in humans. A. Performance over the course of the first training session, measured as % correct averaged over 50-trial moving window. Grey lines, individual participants. Black line, average over participants. B. Performance for best 50-trial window on initial test session after training and (on a subset of participants) upon retesting after a minimum of one week. Each grey dot, one participant; black dots, participants tested on both sessions. Dots are jittered along x axis for visualisation.

### Potential cues for sequence recognition: binary sequences

Which cues are robust correlates of sequence identity, and which can be used by an animal? In our task, possible cues could range across timescales from global to local, as follows (Figure 4A). At the highest (most global) level the GO sequence could be recognised by extracting its overall ordering rule ([3 1 4 2 3 1 4 2] versus other scrambled orders). However, recognition could also stem from detection of specific syllables. For example, in the GO sequence [3 1 4 2 3 1 4 2] the second 100 ms syllable was smaller in magnitude than the first, in contrast to the NO-GO sequence [3 4 2 1 2 4 3 1], whose second syllable was greater than the first (Figure 1D). Thus, detecting a downwards modulation in noise amplitude after 100 ms could enable recognition. Potentially, a more local strategy based on detecting even briefer, sub-syllabic events or “landmarks” in the sequence, such as fluctuations in whisker velocity happening in a certain relative order (Figure 4A), could also be possible. We wondered which strategies were accessible to mice and humans.

**Figure 4.**
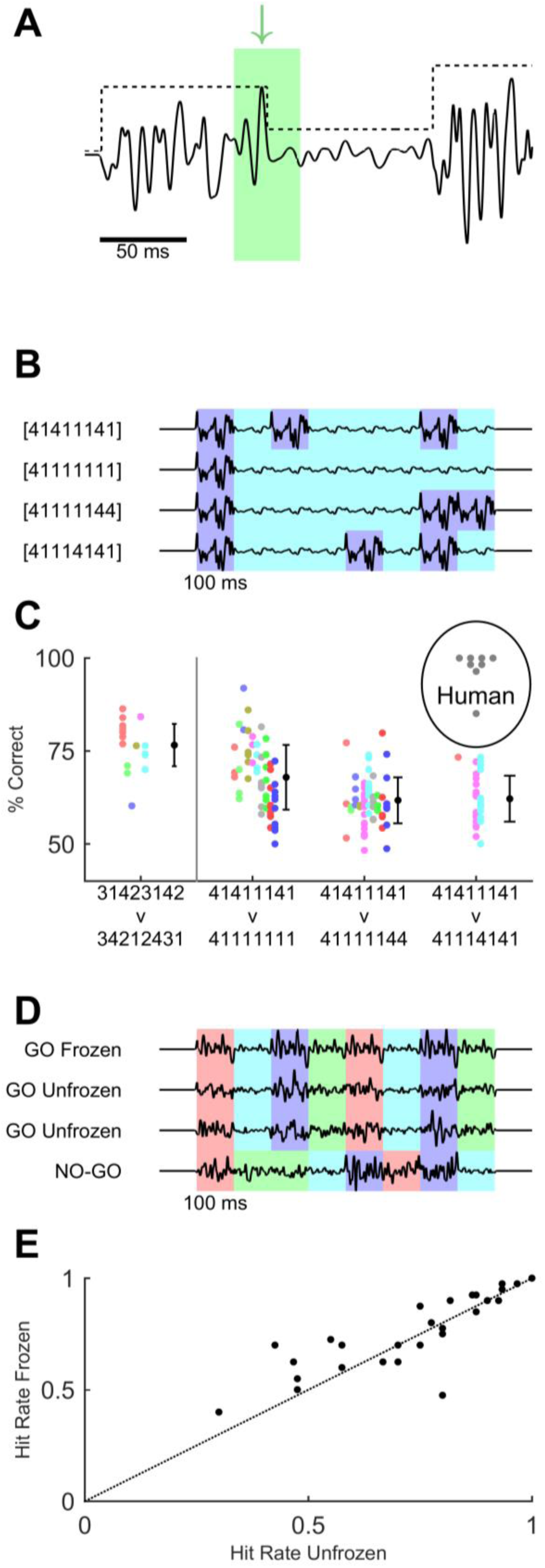
Variations of task design to test for behavioural use of cues. A. Cues within the GO sequence (black line) that could allow recognition of sequence identity. An example of a local cue (within green box) is the large isolated transient “landmark” (green arrow) immediately followed by a low amplitude syllable. In contrast, global cues involve changes in integrated stimulus amplitude over time, as reflected in the amplitude modulation envelope (black dotted line). B. Binary sequences distinguishable based on syllable ordering or the durations of small-amplitude epochs. Different colours indicate epochs of large and small noise amplitude. C. Performance on task variants using binary sequences, compared to original GO vs full NO-GO design (leftmost data point). Each coloured dot indicates a single mouse and session: different mice are in different colours. Black dots and error bars, grand average and SD for each task. Each grey dot indicates performance of an individual human participant on [4 1 4 1 1 1 4 1] vs [4 1 1 1 4 1 4 1]. D. Sequences used to test effect of fixed “landmarks”. Frozen GO used an identical waveform across syllables, trials and sessions. Unfrozen GO maintained the same sequence of noise amplitudes (indicated by colour coding) but varied the detailed waveform across syllables, trials and sessions. NO-GO scrambled the order of syllables, i.e. the sequence of noise amplitudes. E. Hit rate of humans on frozen and unfrozen GO trials. Each dot, one participant and session.

First, any overall temporal average or summation of stimulus parameters throughout the duration of the sequence could be ruled out as a cue, because GO and NO-GO sequences consisted of identical, but scrambled elements. Moreover, animals often started licking on GO trials before the end of sequence presentation (Figure 2D-F), suggesting that just a few transitions in noise modulation or stimulus landmarks sufficed for the animal to reach its decision as to sequence identity.

To begin to explore the ability of mice to exploit specific cues for sequence recognition, we designed a version of the task in which the GO target sequence consisted of a simple succession of epochs of large and small noise amplitude: using the same notation as above, [4 1 4 1 1 1 4 1] (Figure 4B). A separate set of animals (n = 10) was trained to distinguish this new GO sequence from a NO-GO sequence that, as before, differed only in its temporal patterning: [4 1 1 1 4 1 4 1] (Figure 4B). These sequences were simpler than the original design in that they were binary: their constituent syllables were only “large” (4) or “small” (1). These simpler temporal patterns were potentially distinguishable based on syllable ordering or on the different durations of small-amplitude epochs: for example, in the binary GO target sequence the first “small” epoch lasted just 100 ms, but in the NO-GO it lasted for a total of 300 ms (because comprising three syllables). Yet mice performed poorly at distinguishing [4 1 4 1 1 1 4 1] from [4 1 1 1 4 1 4 1] (Figure 4C; n = 3) despite having been trained exclusively on this variant of the task. In particular, animals consistently displayed high false alarm rates, suggesting that they failed to detect what made the binary NO-GO stimulus different from the binary GO (data not shown). This suggested that mice either did not detect the simpler, binary stimulus modulation epochs or did not recognise their differential duration.

To distinguish between these possibilities, we tested performance on probe sessions with two variants of the binary NO-GO sequence. In the first variation, the binary GO target remained identical as [4 1 4 1 1 1 4 1], but the NO-GO sequence was [4 1 1 1 1 1 4 4]. This alternative NO-GO stimulus had the same number of large and small syllables as the one in the previous paragraph, but a different temporal arrangement, with all small-amplitude syllables appearing consecutively and forming a single very long central period (Figure 4B). The sequence therefore effectively had just two “large” epochs, at the beginning and at the end, and a single very long “small” epoch. It was therefore expected to be easier to discriminate from the GO sequence despite having the same overall energy. In the second variation, the binary GO target also remained identical and the NO-GO stimulus was [4 1 1 1 1 1 1 1] (Figure 4B). The goal of this variation was to check whether mice could straightforwardly distinguish large or small syllables. In both of these variants animals performed better than in the original binary design (Figure 4C; p < 10^−9^; n = 7 mice and n = 161 sessions; generalised linear mixed-effects model). Note that performance in the simpler variants was at a level comparable to that of the original GO vs NO-GO paradigm (Figure 4C). Mice also successfully distinguished the binary GO sequence from noise stimuli with no modulation, i.e. with constant noise amplitude: [1 1 1 1 1 1 1 1] or [2 2 2 2 2 2 2 2] (percentage correct > 75% for all animals for [1 1 1 1 1 1 1], 3 out of 4 for [2 2 2 2 2 1 2 2]; n = 4 mice; data not shown). The overall conclusion of the binary sequence experiments is that mice could detect “large” epochs and recognise their number, and use the presence of relative modulations in noise amplitude as cues, but could not as readily use the duration of each modulation epoch.

In contrast to mice, humans rapidly performed well on the [4 1 4 1 1 1 4 1] vs [4 1 1 1 4 1 4 1] version of the task (Figure 4C), even when they had had no prior exposure to the version in Figure 3 (5 out of 8 participants). They achieved high performance within one training session. They also subjectively reported being able to use the relative durations of “small” and “large” epochs as a cue. Thus, humans appeared to have access to more cues for sequence discrimination than mice, including the duration and ordering of intervals in noise modulation.

### Potential cues for sequence recognition: fixed landmarks

Our findings suggested that humans could discriminate sequences based on multiple cues. We wondered whether, in addition to being sensitive to the size and timing of noise amplitude modulations, human participants might also rely on detecting learned sub-syllabic “landmarks”, i.e. specific brief events or fluctuations within the stimulus waveform (Figure 4A). To address this, we assessed whether the presence of fixed waveforms influenced performance.

Upon training participants on the initial version of the task (Figure 3), we tested performance on a variant of the design with two types of GO trials. The first type of trial used a target sequence constructed by applying amplitude modulation to a waveform that was identical (repeated) across syllables and trials (“frozen”). This sequence was used throughout training and in the experiments of Figures 1-3. The second type of trial presented a sequence built by modulating a noise waveform that varied on every repeat (“unfrozen”) (Figure 4D). For unfrozen sequences, each of the 8 syllables was based on a different noise snippet and each trial was constructed from a fresh waveform. Thus, in this type of trial, the amplitude modulation envelope characteristic of the GO sequence remained identical across target trials, but not the precise stimulus values, so that sub-syllabic fluctuations were not conserved. Frozen and unfrozen GO trials were interleaved within a session. Note that unfrozen waveforms could vary in their empirical standard deviation, potentially leading to a confound caused by variability in perceived stimulus amplitude. To control for this, we included in our analysis only stimuli matched for empirical standard deviation. We compared hit rates for both types of GO trial (Figure 4E). Hit rates varied little across type of trial (frozen trials mean 0.76, SD 0.17; unfrozen trials mean 0.73, SD 0.19; p = 0.09; n = 27 participants; Wilcoxon signed rank test). Thus, participants did not require specific brief waveform landmarks to achieve sequence recognition.

In conclusion, humans could use cues based on ordering, timing and feature detection to recognise a target tactile temporal sequence. Mice tested with an identical stimulus paradigm also achieved recognition of a sequence delivered to their whiskers, but appeared to base their performance primarily on the presence of particular relative changes in noise amplitude.

## Discussion

Senses such as touch or hearing depend critically on the detection of temporal patterning over timescales from tens of milliseconds to seconds: in these sensory modalities, signals unfold over time and are incomprehensible if the temporal relationship between their elements is lost. Sequence learning and the processing of temporal duration are impaired in psychiatric disorders including depression and schizophrenia (e.g. [26-29]). Here, we developed an assay suitable for evaluating tactile sequence discrimination. Mice learned to distinguish a target stimulation sequence delivered to their whiskers: the sequence differed from others only in its temporal ordering over hundreds of milliseconds. Humans receiving identical sequential stimuli applied to their fingertip also rapidly learned to perform the task.

Similar human paradigms have been used to discover implicit learning of meaningless auditory noise patterns [30], demonstrating that patterned noise learning generalises across species and sensory systems. In rodents, our design provides an assay for elucidating how neurons within sensory circuits respond and interact under temporally patterned stimulation.

Mechanisms for memorising and determining sequence identity regardless of syntax and semantics have been proposed to be precursors to speech recognition [1, 31, 32]. In our paradigm, sequences were built from chunks of noise with no semantic content or prior meaning. Structural rules such as those aiding interpretation of music or speech (grammar, syntax) were not present as cues. The protocol involved learning only a single instance of a GO target sequence, and did not test generalisation and rule abstraction. However, it would be straightforward to modify the present design to one whereby decisions need to be made based on sequencing or branching rules.

Sequential transitions in texture may be encountered by mice running along walls or tunnels [33, 34] and “receptive” sensation is also routinely mediated by whiskers [23]. Our principal aim in this study was to identify behavioural capacities for learning and recognition of tactile temporal sequences and lay the ground for exploring whether these generalise across domains. A classical approach in comparative cognition employs tasks that are not part of an animal’s natural repertoire to challenge its capacities [35-39].

Although mice and humans were both able to learn to recognise a sequence identifiable by the order of its elements, humans appeared to be capable of accessing a wider range of cues than mice. It is difficult to separate this result from the obvious differences in our ability to communicate task parameters to humans and mice. Our results suggest that mice relied primarily on particular relative fluctuations in stimulus attributes (kinetic or kinematic) over time. Mice could detect “large amplitude” epochs and recognise their number, and could therefore use relative modulations in noise amplitude as cues. In the experiment shown in Figure 2, mice often began licking before the end of the GO target sequence (Figure 2D,F), suggesting that they may have identified the target by detecting particular changes in noise amplitude occurring relatively early in the sequence. In the experiment in Figure 4, mice could not distinguish the GO sequence from others with the same number of “large” and “small” epochs. They did successfully recognise the GO sequence compared to stimuli where the noise amplitude remained constant, regardless of whether the integrated energy of those stimuli matched or exceeded that of the GO sequence, and with no need for prior training (data not shown). This implies that animals used sensitivity to relative changes in noise amplitude as a behavioural cue, a capacity previously demonstrated in rats under a more cognitively demanding task design [40]. Further testing of mouse capacities for abstracting sequencing rules is needed.

Humans appeared to use global and local information on fluctuations in stimulus amplitude to arrive at a heuristic for sequence recognition. Participants receiving the GO sequence often reported feeling a distinct buzzing vibration or counting “beats” in the stimulus, but reported no explicit awareness of a change in sequence element ordering. In a previous auditory study, human listeners were asked to report when a noise stimulus consisted of concatenated repeats of an identical 500 ms segment as opposed to a single 1 s segment [30]. Listeners improved their ability to detect the repeated-noise stimuli when they were unwittingly exposed to the stimulus a few times, and this improvement in performance seemed related to the learning and detection of low-level stimulus waveform features (i.e. particular structures appearing in the noise) [30, 41]. It is likely that the presence of certain learned features, appearing with a specific temporal relationship to each other, provides an elementary cue for discriminating and recognising sequences on timescales of hundreds of milliseconds to seconds across modalities. Determining the duration of the relevant features and how they are encoded [42-44] is a further important task.

Future work must examine how neuronal circuits detect and recognise temporally patterned stimulation sequences. Neurons in early stages of sensory pathways transform any temporally patterned sensory signal into a sequence of precisely timed spikes, so recognising a sensory stimulus with a characteristic temporal pattern –e.g., to discriminate one tactile texture from another [14]– ultimately implies a need for circuits in higher brain areas to decode a spatiotemporal spike sequence. For the paradigm explored here, this capacity is likely to reside within the neocortex, as suggested by the following findings. In the rodent whisker system, neurons in subcortical stages and primary somatosensory cortex display limited temporal integration [45-51]. Therefore, integration over time to represent specific whisker stimulation sequences must be carried out by higher cortical circuits [50-53]. That mice were able to generalise the task across different whisker stimulation directions (Figure 2C), which would have evoked responses in different subsets of neurons at each stage in the pathway [54], suggests further evidence for higher cortical task involvement. A hierarchical scheme whereby later stages of cortical processing can integrate stimuli over longer timescales is consistent with findings in primates [55, 56].

Which mechanisms contribute to setting integration timescales? Single neurons can be sensitive to spatiotemporal input sequences [57-59]. Learning to detect a specific sequence [60] can be accomplished by spike timing-dependent plasticity [61-63]. Timescales for integration of sequences could be regulated by activation of local inhibition [64]. Sequence-selective responses can emerge as a result of sensory exposure to the target [65]. Finally, heterogeneous timescales for integration across cortical processing stages may arise from differences in large-scale connectivity across areas [66, 67]. It remains to be determined how these and other mechanisms come together to implement sequence recognition in cortical circuits in vivo.

## Acknowledgments

This work was supported by the Spanish Ministry of Science and Innovation (grant number BFU2011-23049, co-funded by the European Regional Development Fund; Subprograma Ayudas FPI-MICINN, BES-2012-052293), the Medical Research Council (grant number MR/P006639/1), the Valencia Regional Government (ACOMP2010/199 and PROMETEO/2011/086), and the University of Sussex internal research development fund. The authors declare no competing financial interests.

We thank Karel Svoboda for sharing designs, advice and equipment, Leopoldo Petreanu for sharing designs, Rasmus Petersen for sharing code for behaviour control and for comments on an earlier version of the manuscript, and Elena Giusto for technical help and for the drawings in Figure 1.

## References

1. Wilson, B., Marslen-Wilson, W.D., and Petkov, C.I. (2017). Conserved Sequence Processing in Primate Frontal Cortex. Trends Neurosci 40, 72–82.

2. Lashley, K.S. (1951). The problem of serial order in behavior. In Cerebral Mechanisms in Behavior: The Hixon Symposium, L.A. Jeffress, ed. (New York, NY: John Wiley), pp. 112–146.

3. Griffiths, T.D., and Warren, J.D. (2004). What is an auditory object? Nat Rev Neurosci 5, 887–892.

4. Dehaene, S., Meyniel, F., Wacongne, C., Wang, L., and Pallier, C. (2015). The Neural Representation of Sequences: From Transition Probabilities to Algebraic Patterns and Linguistic Trees. Neuron 88, 2–19.

5. Johnson, K.O. (2001). The roles and functions of cutaneous mechanoreceptors. Curr Opin Neurobiol 11, 455–461.

6. Johansson, R.S., and Birznieks, I. (2004). First spikes in ensembles of human tactile afferents code complex spatial fingertip events. Nat Neurosci 7, 170–177.

7. Bale, M.R., Campagner, D., Erskine, A., and Petersen, R.S. (2015). Microsecond-scale timing precision in rodent trigeminal primary afferents. J Neurosci 35, 5935–5940.

8. Chagas, A.M., Theis, L., Sengupta, B., Stuttgen, M.C., Bethge, M., and Schwarz, C. (2013). Functional analysis of ultra high information rates conveyed by rat vibrissal primary afferents. Front Neural Circuits 7, 190.

9. Jones, L.M., Lee, S., Trageser, J.C., Simons, D.J., and Keller, A. (2004). Precise temporal responses in whisker trigeminal neurons. J Neurophysiol 92, 665–668.

10. Campagner, D., Evans, M.H., Bale, M.R., Erskine, A., and Petersen, R.S. (2016). Prediction of primary somatosensory neuron activity during active tactile exploration. Elife 5.

11. Bush, N.E., Schroeder, C.L., Hobbs, J.A., Yang, A.E., Huet, L.A., Solla, S.A., and Hartmann, M.J. (2016). Decoupling kinematics and mechanics reveals coding properties of trigeminal ganglion neurons in the rat vibrissal system. Elife 5.

12. Phillips, J.R., and Johnson, K.O. (1985). Neural mechanisms of scanned and stationary touch. The Journal of the Acoustical Society of America 77, 220–224.

13. Phillips, J.R., Johansson, R.S., and Johnson, K.O. (1990). Representation of braille characters in human nerve fibres. Exp Brain Res 81, 589–592.

14. Weber, A.I., Saal, H.P., Lieber, J.D., Cheng, J.W., Manfredi, L.R., Dammann, J.F., 3rd, and Bensmaia, S.J. (2013). Spatial and temporal codes mediate the tactile perception of natural textures. Proc Natl Acad Sci U S A 110, 17107–17112.

15. Saal, H.P., Wang, X., and Bensmaia, S.J. (2016). Importance of spike timing in touch: an analogy with hearing? Curr Opin Neurobiol 40, 142–149.

16. Maravall, M., and Diamond, M.E. (2014). Algorithms of whisker-mediated touch perception. Curr Opin Neurobiol 25, 176–186.

17. Sofroniew, N.J., and Svoboda, K. (2015). Whisking. Curr Biol 25, R137–140.

18. Guo, Z.V., Hires, S.A., Li, N., O’Connor, D.H., Komiyama, T., Ophir, E., Huber, D., Bonardi, C., Morandell, K., Gutnisky, D., et al. (2014). Procedures for behavioral experiments in head-fixed mice. PLoS One 9, e88678.

19. O’Connor, D.H., Clack, N.G., Huber, D., Komiyama, T., Myers, E.W., and Svoboda, K. (2010). Vibrissa-based object localization in head-fixed mice. J Neurosci 30, 1947–1967.

20. Hayar, A., Bryant, J.L., Boughter, J.D., and Heck, D.H. (2006). A low-cost solution to measure mouse licking in an electrophysiological setup with a standard analog-to-digital converter. J Neurosci Methods 153, 203–207.

21. Tucci, V., Hardy, A., and Nolan, P.M. (2006). A comparison of physiological and behavioural parameters in C57BL/6J mice undergoing food or water restriction regimes. Behav Brain Res 173, 22–29.

22. Carandini, M., and Churchland, A.K. (2013). Probing perceptual decisions in rodents. Nat Neurosci 16, 824–831.

23. Diamond, M.E., and Arabzadeh, E. (2013). Whisker sensory system - from receptor to decision. Prog Neurobiol 103, 28–40.

24. Green, D.M., and Swets, J.A. (1966). Signal Detection Theory and Psychophysics, (New York: Wiley).

25. Berditchevskaia, A., Caze, R.D., and Schultz, S.R. (2016). Performance in a GO/NOGO perceptual task reflects a balance between impulsive and instrumental components of behaviour. Sci Rep 6, 27389.

26. Fischer, F. (1929). Zeitstruktur und Schizophrenie. Zeitschrift für die gesamte Neurologie und Psychiatrie 121, 544–574.

27. Marvel, C.L., Schwartz, B.L., Howard, D.V., and Howard, J.H., Jr. (2005). Implicit learning of non-spatial sequences in schizophrenia. J Int Neuropsychol Soc 11, 659–667.

28. Siegert, R.J., Weatherall, M., and Bell, E.M. (2008). Is implicit sequence learning impaired in schizophrenia? A meta-analysis. Brain Cogn 67, 351–359.

29. Abrahamse, E.L., van der Lubbe, R.H., and Verwey, W.B. (2009). Sensory information in perceptual-motor sequence learning: visual and/or tactile stimuli. Exp Brain Res 197, 175–183.

30. Agus, T.R., Thorpe, S.J., and Pressnitzer, D. (2010). Rapid formation of robust auditory memories: insights from noise. Neuron 66, 610–618.

31. Petkov, C.I., and Jarvis, E.D. (2012). Birds, primates, and spoken language origins: behavioral phenotypes and neurobiological substrates. Front Evol Neurosci 4, 12.

32. Comins, J.A., and Gentner, T.Q. (2014). Temporal pattern processing in songbirds. Curr Opin Neurobiol 28, 179–187.

33. Jenks, R.A., Vaziri, A., Boloori, A.R., and Stanley, G.B. (2010). Self-motion and the shaping of sensory signals. J Neurophysiol 103, 2195–2207.

34. Sofroniew, N.J., Cohen, J.D., Lee, A.K., and Svoboda, K. (2014). Natural whisker-guided behavior by head-fixed mice in tactile virtual reality. J Neurosci 34, 9537–9550.

35. Gould, J.L. (2004). Animal cognition. Curr Biol 14, R372–375.

36. Roth, G., and Dicke, U. (2005). Evolution of the brain and intelligence. Trends Cogn Sci 9, 250–257.

37. Alem, S., Perry, C.J., Zhu, X., Loukola, O.J., Ingraham, T., Sovik, E., and Chittka, L. (2016). Associative Mechanisms Allow for Social Learning and Cultural Transmission of String Pulling in an Insect. PLoS Biol 14, e1002564.

38. Loukola, O.J., Perry, C.J., Coscos, L., and Chittka, L. (2017). Bumblebees show cognitive flexibility by improving on an observed complex behavior. Science 355, 833–836.

39. Ishiyama, S., and Brecht, M. (2016). Neural correlates of ticklishness in the rat somatosensory cortex. Science 354, 757–760.

40. Fassihi, A., Akrami, A., Esmaeili, V., and Diamond, M.E. (2014). Tactile perception and working memory in rats and humans. Proc Natl Acad Sci U S A 111, 2331–2336.

41. Andrillon, T., Kouider, S., Agus, T., and Pressnitzer, D. (2015). Perceptual learning of acoustic noise generates memory-evoked potentials. Curr Biol 25, 2823–2829.

42. Jadhav, S.P., Wolfe, J., and Feldman, D.E. (2009). Sparse temporal coding of elementary tactile features during active whisker sensation. Nat Neurosci 12, 792–800.

43. Waiblinger, C., Brugger, D., and Schwarz, C. (2015). Vibrotactile discrimination in the rat whisker system is based on neuronal coding of instantaneous kinematic cues. Cereb Cortex 25, 1093–1106.

44. Waiblinger, C., Brugger, D., Whitmire, C.J., Stanley, G.B., and Schwarz, C. (2015). Support for the slip hypothesis from whisker-related tactile perception of rats in a noisy environment. Front Integr Neurosci 9, 53.

45. Maravall, M., Petersen, R.S., Fairhall, A.L., Arabzadeh, E., and Diamond, M.E. (2007). Shifts in coding properties and maintenance of information transmission during adaptation in barrel cortex. PLoS Biol 5, e19.

46. Jacob, V., Le Cam, J., Ego-Stengel, V., and Shulz, D.E. (2008). Emergent properties of tactile scenes selectively activate barrel cortex neurons. Neuron 60, 1112–1125.

47. Petersen, R.S., Brambilla, M., Bale, M.R., Alenda, A., Panzeri, S., Montemurro, M.A., and Maravall, M. (2008). Diverse and temporally precise kinetic feature selectivity in the VPm thalamic nucleus. Neuron 60, 890–903.

48. Stuttgen, M.C., and Schwarz, C. (2010). Integration of vibrotactile signals for whisker-related perception in rats is governed by short time constants: comparison of neurometric and psychometric detection performance. J Neurosci 30, 2060–2069.

49. Estebanez, L., El Boustani, S., Destexhe, A., and Shulz, D.E. (2012). Correlated input reveals coexisting coding schemes in a sensory cortex. Nat Neurosci 15, 1691–1699.

50. Pitas, A., Albarracin, A.L., Molano-Mazon, M., and Maravall, M. (2016). Variable Temporal Integration of Stimulus Patterns in the Mouse Barrel Cortex. Cereb Cortex.

51. McGuire, L.M., Telian, G., Laboy-Juarez, K.J., Miyashita, T., Lee, D.J., Smith, K.A., and Feldman, D.E. (2016). Short Time-Scale Sensory Coding in S1 during Discrimination of Whisker Vibrotactile Sequences. PLoS Biol 14, e1002549.

52. Lim, Y., Lagoy, R., Shinn-Cunningham, B.G., and Gardner, T.J. (2016). Transformation of temporal sequences in the zebra finch auditory system. Elife 5.

53. Fassihi, A., Akrami, A., Schönfelder, V.H., and Diamond, M.E. (2015). Temporal integration in a vibrotactile delayed comparison task: From sensory coding to decision in humans and rats. In Society for Neuroscience Meeting. (Chicago, IL).

54. Bale, M.R., and Petersen, R.S. (2009). Transformation in the neural code for whisker deflection direction along the lemniscal pathway. J Neurophysiol 102, 2771–2780.

55. Hasson, U., Yang, E., Vallines, I., Heeger, D.J., and Rubin, N. (2008). A hierarchy of temporal receptive windows in human cortex. J Neurosci 28, 2539–2550.

56. Honey, C.J., Thesen, T., Donner, T.H., Silbert, L.J., Carlson, C.E., Devinsky, O., Doyle, W.K., Rubin, N., Heeger, D.J., and Hasson, U. (2012). Slow cortical dynamics and the accumulation of information over long timescales. Neuron 76, 423–434.

57. Segundo, J.P., Moore, G.P., Stensaas, L.J., and Bullock, T.H. (1963). Sensitivity of Neurones in Aplysia to Temporal Pattern of Arriving Impulses. The Journal of experimental biology 40, 643–667.

58. Branco, T., Clark, B.A., and Hausser, M. (2010). Dendritic discrimination of temporal input sequences in cortical neurons. Science 329, 1671–1675.

59. Baker, C.A., Ma, L., Casareale, C.R., and Carlson, B.A. (2016). Behavioral and Single-Neuron Sensitivity to Millisecond Variations in Temporally Patterned Communication Signals. J Neurosci 36, 8985–9000.

60. Hardy, N.F., and Buonomano, D.V. (2016). Neurocomputational Models of Interval and Pattern Timing. Curr Opin Behav Sci 8, 250–257.

61. Masquelier, T., Guyonneau, R., and Thorpe, S.J. (2008). Spike timing dependent plasticity finds the start of repeating patterns in continuous spike trains. PLoS One 3, e1377.

62. Klampfl, S., and Maass, W. (2013). Emergence of dynamic memory traces in cortical microcircuit models through STDP. J Neurosci 33, 11515–11529.

63. Gütig, R. (2014). To spike, or when to spike? Curr Opin Neurobiol 25, 134–139.

64. Kepecs, A., and Fishell, G. (2014). Interneuron cell types are fit to function. Nature 505, 318–326.

65. Gavornik, J.P., and Bear, M.F. (2014). Learned spatiotemporal sequence recognition and prediction in primary visual cortex. Nat Neurosci 17, 732–737.

66. Chaudhuri, R., Bernacchia, A., and Wang, X.J. (2014). A diversity of localized timescales in network activity. Elife 3, e01239.

67. Chaudhuri, R., Knoblauch, K., Gariel, M.A., Kennedy, H., and Wang, X.J. (2015). A Large-Scale Circuit Mechanism for Hierarchical Dynamical Processing in the Primate Cortex. Neuron 88, 419–431.

